# Human episodic memory retrieval is accompanied by a neural contiguity effect

**DOI:** 10.1101/117010

**Authors:** Sarah Folkerts, Ueli Rutishauser, Marc W. Howard

## Abstract

Cognitive psychologists have long hypothesized that experiences are encoded in a temporal context that changes gradually over time. When an episodic memory is retrieved, the state of context is recovered—a jump back in time. We recorded from single units in the MTL of epilepsy patients performing an item recognition task. The population vector changed gradually over minutes during presentation of the list. When a probe from the list was remembered with high confidence, the population vector reinstated the temporal context of the original presentation of that probe during study—a neural contiguity effect that provides a possible mechanism for behavioral contiguity effects. This pattern was only observed for well-remembered probes; old probes that were not well-remembered showed an anti-contiguity effect. These results constitute the first direct evidence that recovery of an episodic memory in humans is associated with retrieval of a gradually-changing state of temporal context—a neural “jump-back-in-time” that parallels the act of remembering.

**Significance statement:** Episodic memory is the ability to re-live a specific experience from one’s life. For decades, researchers have hypothesized that, unlike other forms of memory that can be described as simple associations between stimuli, episodic memory depends on the recovery of a neural representation of spatiotemporal context. During study of a sequence of stimuli, the brain state of epilepsy patients changed slowly over at least a minute. When the participant remembered a particular event from the list, this gradually-changing state was recovered. This provides direct confirmation of the prediction from computational models of episodic memory. The resolution of this point means that the study of episodic memory can focus on the mechanisms by which this representation of spatiotemporal context is maintained and, sometimes recovered.

## Introduction

Episodic memory refers to our ability to vividly remember specific events from our own experience. The vividness of episodic memory, along with the specificity of the memory to a particular place and time has led researchers to characterize episodic memory as mental time travel (Tulving, 1972; Hassabis, Kumaran, Vann, & Maguire, 2007; Schacter, Addis, & Buckner, 2007). This verbal description has been operationalized in computational models of episodic memory in which the flow of time is described by a slowly and gradually-changing state of temporal context (Howard & Kahana, 2002; Sederberg, Howard, & Kahana, 2008; Polyn, Norman, & Kahana, 2009; Howard, Shankar, Aue, & Criss, 2015). In these models episodic memory retrieval is accompanied by the recovery of a prior state of temporal context—a jump back in time—that accounts for the behavioral contiguity effect (Kahana, 1996; Schwartz, Howard, Jing, & Kahana, 2005; Howard, Youker, & Venkatadass, 2008; Unsworth, 2008).

This computational hypothesis makes two predictions that can be tested neurally (see Fig. 1). First, in addition to stimulus-evoked activity, the activity of some neurons involved in episodic memory should also change gradually over time. This prediction aligns with a large body of animal work showing that neural ensembles in the hippocampus, amygdala and prefrontal cortex change slowly over time scales up to at least tens of minutes (Manns, Howard, & Eichenbaum, 2007; Hyman, Ma, Balaguer-Ballester, Durstewitz, & Seamans, 2012; Mankin et al., 2012; MacDonald, Lepage, Eden, & Eichenbaum, 2011; Salz et al., 2016; Rubin, Geva, Sheintuch, & Ziv, 2015; Cai et al., 2016; Rashid et al., 2016; Howard, 2017). Second, during retrieval of an existing memory, the prior state (temporal context) associated with an episodic memory should be restored. Although some prior studies have attempted to measure this hypothesized reinstatement (Manning, Polyn, Litt, Baltuch, & Kahana, 2011; Howard, Viskontas, Shankar, & Fried, 2012; Yaffe et al., 2014), due to methodological limitations of those studies there is presently no definitive study linking recovery of a gradually-changing temporal context to episodic memory in humans.

**Figure 1.**
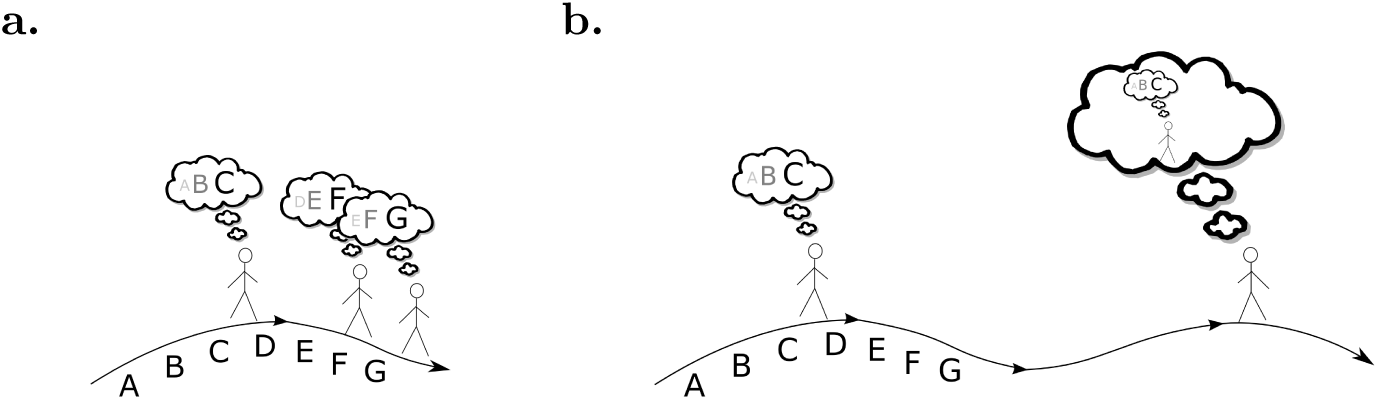
A neural signature of retrieved temporal context. **a.** While experiencing a sequence of stimuli A B C … the brain is hypothesized to maintain information about the recent past at each moment. Because the recent past changes gradually, so too should this brain state. That is the brain state after G should resemble the brain state after F moreso than the brain state after C. This gradually-changing representation is hypothesized to form a temporal context for the study items. **b.** Retrieved temporal context models hypothesize that an episodic memory is accompanied by recovery of the temporal context at the time that memory was encoded. When the participant remembers a particular event such as C, this reinstates the temporal context when C was experienced. This predicts that the brain state after memory for C should resemble the brain state during experience of the *neighbors* of **c.** The similarity should fall off with distance from C in both the forward and backward directions.

Episodic memory is often studied in the laboratory using the item recognition task. In item recognition, participants are presented with a study list of novel stimuli to remember (here we use pictures). After study, participants are provided with a set of probe stimuli one at a time, some of which were on the study list and some of which were not. The participants’ task is to distinguish probe stimuli that were on the list from probe stimuli that were not on the list. Many authors have hypothesized that recognition memory is supported by two processes, recollection and familiarity (Yonelinas et al., 2002; Eichenbaum, Yonelinas, & Ranganath, 2007; Staresina, Fell, Dunn, Axmacher, & Henson, 2013; Wixted, 2007). According to this viewpoint (which it should be noted is not universally accepted, (Squire, Wixted, & Clark, 2007)) recollection corresponds to vivid episodic memory in which details of the study experience is recovered. When an old probe is recollected, triggering retrieval of an episodic memory, this is believed to lead participants to endorse the probe as old with high confidence (Yonelinas et al., 2002; Diana, Yonelinas, & Ranganath, 2007). Regardless of one’s position on two-process theory, it is clear that highest confidence old responses are often associated with the recovery of detailed source information about the context in which a probe was studied (Onyper, Zhang, & Howard, 2010; Slotnick & Dodson, 2005; Hautus, Macmillan, & Rotello, 2008), with a behavioral contiguity effect (Schwartz et al., 2005), and with the activation of neurons in the medial temporal lobe (Rutishauser et al., 2015), properties that we would ordinarily associate with an episodic memory. In this study we will operationalize highest confidence old responses as a marker of probes that were more likely to have triggered an episodic memory.

## Methods and Materials

In this study epilepsy patients performed an item recognition task, rating their confidence that probes were presented on a six-point scale (Figure 2). During both study and retrieval, single units were recorded from microelectrodes implanted in their medial temporal lobes. Population vectors were measured across units; consistent with previous results the population vectors changed gradually during study of the list. Comparing the population vector in response to an old probe at test to the population vectors during study will enable us to evaluate whether temporal context is recovered. We test the hypothesis that probes that triggered a strong episodic memory—here operationalized as probes that received a highest-confidence response—are accompanied by greater recovery of temporal context than probes that did not trigger a strong episodic memory—here operationalized as probes that did not receive a highest-confidence response.

**Figure 2.**
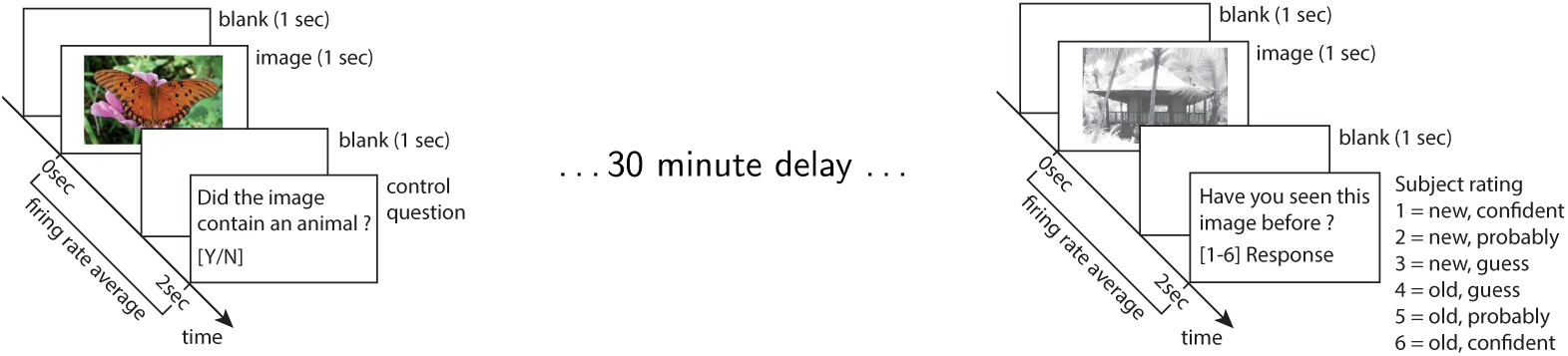
The behavioral task. During a study (learning) phase, participants were asked to learn set of pictures. In order to ensure that the patients were attending to the picture, they responded to an orienting task after each item. After a 30 minute delay, participants were presented with a test list that included both stimuli from the study session and also new probes. For each, they indicated whether they thought they had seen an item before or not on a 6-point confidence scale.

### Patients

54 recording sessions were made from 35 patients of either sex who were evaluated for possible surgical treatment of epilepsy using implantation of depth electrodes. All patients volunteered for the study and gave informed consent. Protocols were approved by the institutional review boards of the Cedars-Sinai Medical Center, Huntington Memorial Hospital and the California Institute of Technology. Out of the 54 recording sessions, 44 were previously reported by Rutishauser et al. (2015) and 10 were not. The dataset used in this paper is a subset of a publicly available dataset that has been published along with a paper that describes the methods in detail (Faraut et al., 2018b, 2018a).^1^ Five sessions were rejected because memory accuracy was not sufficiently high (*d*^′^ < .5). The remaining 49 sessions were from 33 patients, of whom 24 were male and 9 were female.

### Electrophysiology and spike sorting

The recording methods and single-unit data analyses for this dataset have been described in detail before (Rutishauser et al., 2015; Faraut et al., 2018b). Briefly, the recordings analyzed here were obtained from depth electrodes implanted bilaterally within the hippocampus and amygdala (8 microwires each, 32 channels per patient in total). Broadband extracellular recordings were filtered .1 Hz to 9 kHz and sampled at 32 kHz (Neuralynx Inc). Electrodes were localized based on post-operative MRI images. Electrode locations were chosen according to clinical criteria alone. Spikes were detected and sorted as previously described (Rutishauser, Mamelak, & Schuman, 2006).

### Behavioral task

The task (Figure 2) consisted of two parts: a study (learning) phase followed by a test phase. During study, patients viewed a list of 100 photographs of natural scenes. There were 25 instances each from five different visual categories (animals, people, cars/vehicles, outdoor scenes/houses and flowers/food items; see Figure 6 for examples). The list was randomly assembled such that categories were not clustered. Each image appeared on the screen for 1 s, followed by a blank delay of 0.5 s, followed by an orienting task in which participants answered whether the image they had just seen contained an animal or not. The method used in this behavioral task is the same as that used in Rutishauser et al. (2015)

A delay that ranged in duration from about 15 minutes to about 30 minutes intervened between study of the last stimulus and the beginning of the test list. During the test phase, subjects were shown 100 images, half of which were identical to those seen previously (“old”) and half were novel (“new”). After each image, subjects indicated whether they saw the image before or not together with how confident they were about their decision (1= “new, certain”, 2 = “new, probably”, 3 = “new, guess”, 4 = “old, guess”, 5 = “old, probably”, 6 = “old, certain”). There was no response deadline.

### Artifact rejection

We excluded 96 units that contributed no spikes to the firing rate vectors or that had a bimodal firing rate distribution. This left a total number of 1286 units used in this report. In addition, we excluded trials during which there was an abrupt signal loss in several simultaneously recorded units (Figure 3). Such loss is likely attributable to recording problems and we thus excluded such periods of time. To achieve this, time periods during which a fraction of ≥ .25 of the units ceased firing for ≥ .05 of the total trial duration were classified as artifacts. We identified such artifacts in two study and four test sessions.

**Figure 3.**
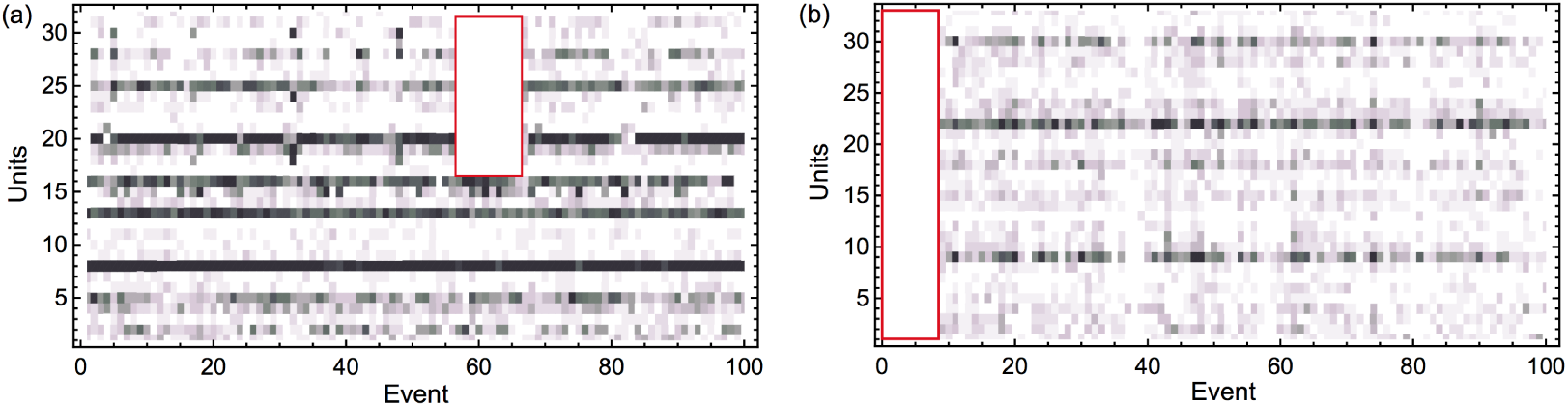
Artifact rejection. The raster plots show the activity of each unit (row) as a function of time. The method for artifact rejection described in the text identified the red squares as an artifact. The rejected units were all located in the same brain region.

### Population vectors

Population vectors were computed from the average firing rates within a 2 s window starting at stimulus onset. To control for changes in baseline firing rate for different units, the mean firing rates for each unit were z-scored with respect to the average firing rate of that unit across all events. After z-scoring, all statistics reported were computed across all recorded units across all sessions. Trials with reaction times that exceeded 2.5 standard deviations of the reaction time distribution of a given patient were excluded (136/10439 trials were excluded based on this criterion).

### Recency analysis

In order to evaluate whether the ensemble changed gradually over time we analyzed how the similarity between population vectors changed as a function of recency, the difference between the serial position of the two events. For instance, the comparison between the population vector from presentation of the seventh stimulus in the list to the population vector from the fourth stimulus in the list is associated with a recency of −3 (see Figure 4a). A normalized inner product of z-scored population vectors (the inner product normalized by the number of units) was used to characterise the similarity between the ensemble response between a pair of events as a function of recency. In order to avoid any possible confounding influence of a primacy effect on the analysis, only events after the first 20 item presentations were included. In doing statistics on effects of recency and contiguity (described below), recording session was treated as a random variable.

**Figure 4.**
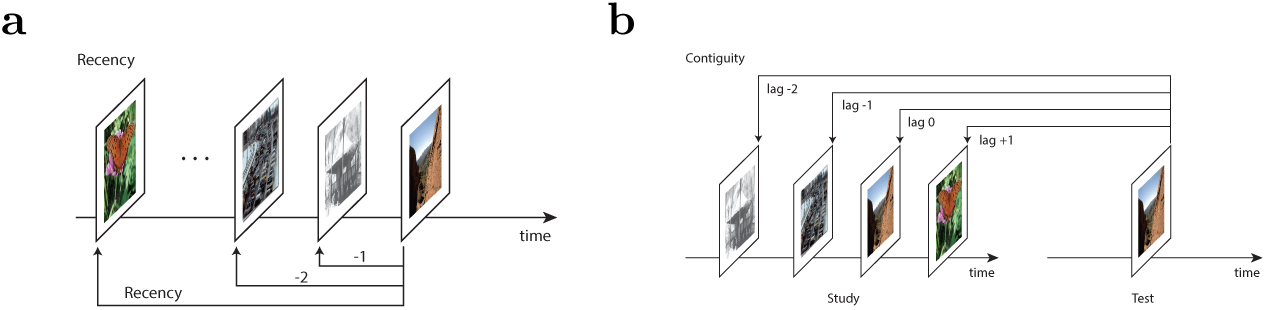
Schematic for definition of recency and contiguity. Analyses in this paper compute the neural pattern similarity between pairs of events. These similarities are averaged over the experiment and aggregated as a function of recency or contiguity. **a.** Recency is defined as the difference in the serial positions at which two events took place. **b.** Contiguity is measured in units of lag. When a stimulus is presented as a recognition probe, lag is defined as the difference in serial positions between the original presentation of the probe stimulus. Comparison of a probe to the original presentation of that stimulus is associated with a lag of zero.

### Neural contiguity analyses

In order to evaluate whether memory for an event caused reconstruction of the gradually-changing neural state during study of that event we compared ensemble similarity as a function of lag, defined as follows (see Figure 4b). Given an old test probe that was originally (during study) presented at serial position *i*, and a study event presented at serial position *j*, lag is defined as *j−i*. To be concrete, consider the population vector from the test of an old probe originally presented at serial position seven. We compared this population vector to each of the population vectors from a study event. The lag associated with comparison of the test event to the study event at serial position seven—the same stimulus—is zero. The lag associated with the comparison to the population vector from study at serial position eight—which immediately followed study of the stimulus—is +1; the lag associated with the comparison to the event from serial position six is −1. For each old probe, lag defines a number line across the study serial positions with zero at the original study location of the probe stimulus. Ensemble similarity between each pair of events consisting of an old test probe and a study stimulus was aggregated as a function of lag.

Note that the number of data points entering into the contiguity analyses changes as a function of lag. For instance, there are many more combinations of serial positions that result in a lag of +1 than there are combinations that lead to a lag of +50. For statistical tests, we restricted our attention to lags between −30 and +30. Because lag zero is a special case (similarity could be boosted simply because the same visual features present during study and test of the same stimulus), lag zero was not included in statistical analyses of lag.

### Isolating a neural signature of episodic memory—memory advantage index

A goal of this work is to identify the neural correlates of episodic memory. Here, we used confidence ratings to compare between memories with large *vs* small episodic contributions. A large body of work has argued that responses to old items that do not receive the highest confidence response rely on familiarity (and perhaps weak recollection), whereas highest-confidence old responses rely on a mixture of familiarity and strong recollection (Yonelinas et al., 2002; Diana et al., 2007). In order to isolate the contribution due to episodic memory, we computed a difference between old probes that received a highest-confidence response and old probes that did not receive a highest-confidence response. ^2^ Note that this analysis can only identify neural signatures of episodic memory performance that manifest as consistent changes as a function of temporal lag—there may well be other neural signatures of episodic memory that are invisible to this analysis.

In order to compute this difference due to memory we started by taking the product of z-scored firing rates for each pair of stimuli that entered into the contiguity analysis aggregated by lag. However, rather than averaging over units, as in taking the normalized inner product we computed a matrix with each possible lag corresponding to the columns and each unit corresponding to the rows. Separate matrices were computed using the similarity for low-confidence and high confidence trials. To estimate a difference attributable to episodic memory, we took a paired t-test (over units) for each lag as a measure of “memory advantage”. The use of the paired t-statistic minimizes variability due to difference in the units.

The t-statistic can be used to evaluate the null hypothesis directly (values greater than 1.96 are statistically different from zero), but also the t-statistic can be compared for different lags. If there was no recovery of temporal context, the memory advantage would be the same across lags; a systematic change in the memory advantage as a function of lag must reflect recovery of some form of information that was present during study of the list.

### Permutation analysis

The assumptions of the traditional parametric tests used in the contiguity analysis are violated. For instance, z-scored firing rates are in general non-normal. In order to eliminate concerns that the conclusions were simply the result of inappropriate parametric statistics, we supplemented those analyses with a permutation test. In this permutation analysis the stimulus identities during the test session were randomly shuffled thereby removing any actual link to the actual study events. We separately permuted the identity of all of the old probes that were remembered with highest confidence (6) and all other confidences (1-5) independently among themselves. We then recomputed the statistics reported in the contiguity analysis for each of 1000 random permutations. This procedure preserves all of the marginal distributions. However, it disrupts the actual temporal relationships between study and test. If the observed effect of a temporal variable (e.g., |lag*|*) exceeds the distribution of the permuted data, this supports the conclusions of the parametric statistics.

### Separate contiguity analyses for hippocampus and amygdala

In order to gain further insight into the anatomical origins of the contiguity signal we examined contiguity effects separately for gross anatomical regions. We computed contiguity effects separately for units recorded from the amygdala and hippocampus, collapsing over hemispheres. Note that there are more units recorded from the amygdala (849) compared to the hippocampus (533).

### Additional analyses of lower-confidence responses

We also conducted an analogous set of analyses in which we compared three types of responses: highest confidence old responses (6), lower confidence old responses (4-5), and misses, old probes that received a new response (1-3).

### Visually-selective (VS) units

In order to determine if the gradually-changing temporal context representations examined using the recency and contiguity analyses were distinct from visual representations we repeated these analyses restricting the analysis to visually-selective (VS) units. VS units are those that responded differently to the different categories of images, as assessed by an ANOVA on their firing rate. Methods for identifying VS units were identical to those reported in detail previously (Rutishauser et al., 2015). 213/1286 units were classified as visually-selective.

## Results

### Behavioral results were consistent with episodic memory for some old probes

Patients judged each item presented during the test phase as either old (seen before) or new (not seen before) together with a confidence rating (Figure 2). The behavioral results from patients were broadly consistent with canonical behavioral results from control participants (Kahana, 2012). Patients used all confidence ratings and used the highest confidence old response approximately five times more often for old probes than for new probes (Figure 5a). In contrast, the lower confidence old responses (4-5) were less effective in discriminating old probes from new probes. Patients were able to differentiate weak from strong memories using subjective confidence ratings.

**Figure 5.**
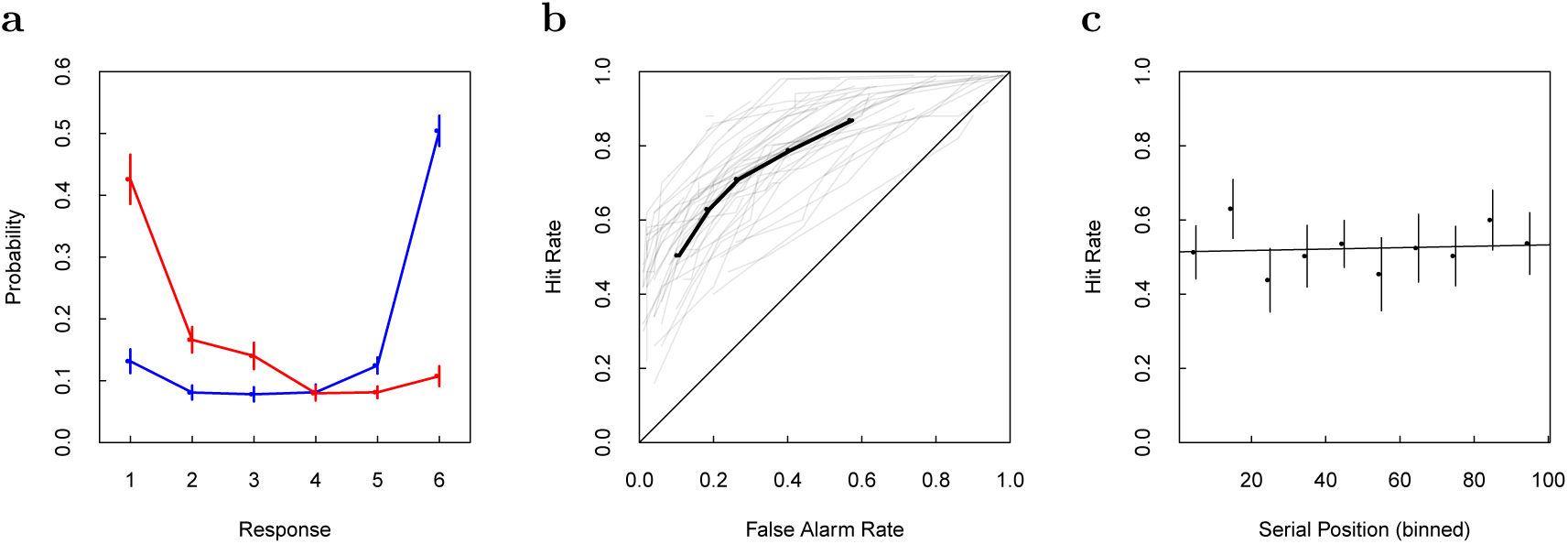
Behavioral results. **a.** Participants successfully distinguished repeated probes from new probes. Shown is the probability of each response (1-6) conditional on the ground truth, i.e. whether the stimulus is old (blue) or new (red). Note that responses (1,2,3) for new (red) stimuli and responses (4,5,6) for old (blue) stimuli are correct whereas the others are incorrect. Patients had good memory as demonstrated by using the highest confidence rating (1 or 6) for about half of the new and old probes, respectively. Error bars are SEM across n=49 sessions. **b.** Behavioral ROC curves for each participant included in this study (grey lines) and the average ROC (red dashed line). The ROC plots hit rate as a function of false alarm rate for each possible criterion; chance performance would be along the diagonal. These ROC curves are typical of item recognition studies, with a reliable asymmetry characteristic of episodic memories (see text for details). **c.** The 30 minute delay between study and test successfully eliminated behavioral recency effect. The hit rate—here the probability of an old probe receiving a highest-confidence response—is shown as a function of each probe’s binned serial position during study. The slope of the regression line is not significantly different from zero. Error bars are the 95% confidence interval.

We next quantified each patient’s behavior using a receiver operating characteristics (ROC) curve (Figure 5b). The ROC shows hit rate—probability of a yes response to an *old* probe—as a function of false alarm rate—probability of a yes response to a *new* probe—for each possible confidence criterion. To the extent the ROC points lie above the diagonal, memory is above chance. The ROC curves were asymmetric (Figure. 5b). To quantify this we computed the slope of the z-transformed ROC curve for each session and compared the values to one. The average slope .73 ± .04, mean *±* SEM, was reliably less than one, *t*(46) = 6.32, *p* < .001 and the slope was less than one for 40 out of 49 sessions.^3^

A slope of the z-ROC curve less than one is commonly taken as a signature of episodic memory accompanied by successful recollection (Yonelinas, 2002; Wixted, 2007; Fortin, Wright, & Eichenbaum, 2004).

It is important for the later neural analysis that the probability of recognizing a stimulus as old does not depend on how long ago it was seen; some previous attempts to measure a neural jump-back-in-time (Howard et al., 2012), were confounded by a large behavioral recency effect. In this study, the study and test period were separated by a delay of approximately 30 min. This successfully eliminated the recency effect: the hit rate (probability of a yes response) was independent of the position an item was shown in during study (Figure 5c). There was no significant effect of serial position, as indicated by a regression coefficient, (2 *±* 7) *×* 10^−4^, that was not reliably different from zero. This shows that the delay between study and test was effective in eliminating the recency effect at test. These behavioral results are consistent with another study using the same task (Figure 1, Rutishauser, Aflalo, Rosario, Pouratian, & Andersen, 2017).

### The population vector during study changed gradually over at least a minute

A key requirement for contextual reinstatement to occur is that neural activity changes gradually across multiple stimulus presentations during learning. Because they are not imposed by the stimuli, which are randomized in order, such gradual changes could be a signature of temporal information. We thus first tested whether neurons within the MTL exhibit signatures of a gradually changing temporal context. We constructed population vectors from the mean firing rate in a 2 s window following stimulus onset for all recorded units (see methods for details). We then tested whether the pairwise similarity between population vectors from study events differed systematically as a function of time between those events. We found a gradual increase in similarity for pairs of study events closer together in time (Figure 6a); the regression coefficient was .00123 ± .00008, *F*_(1,78)_ = 221.6, *p* < .001. A similar recency effect was also evident during the test phase (Figure 9); the regression coefficient was .00089 ±.00006, *F*_(1,78)_ = 188.7, *p* < .001. We thus found significant temporal context effects during both study and test.

**Figure 6.**
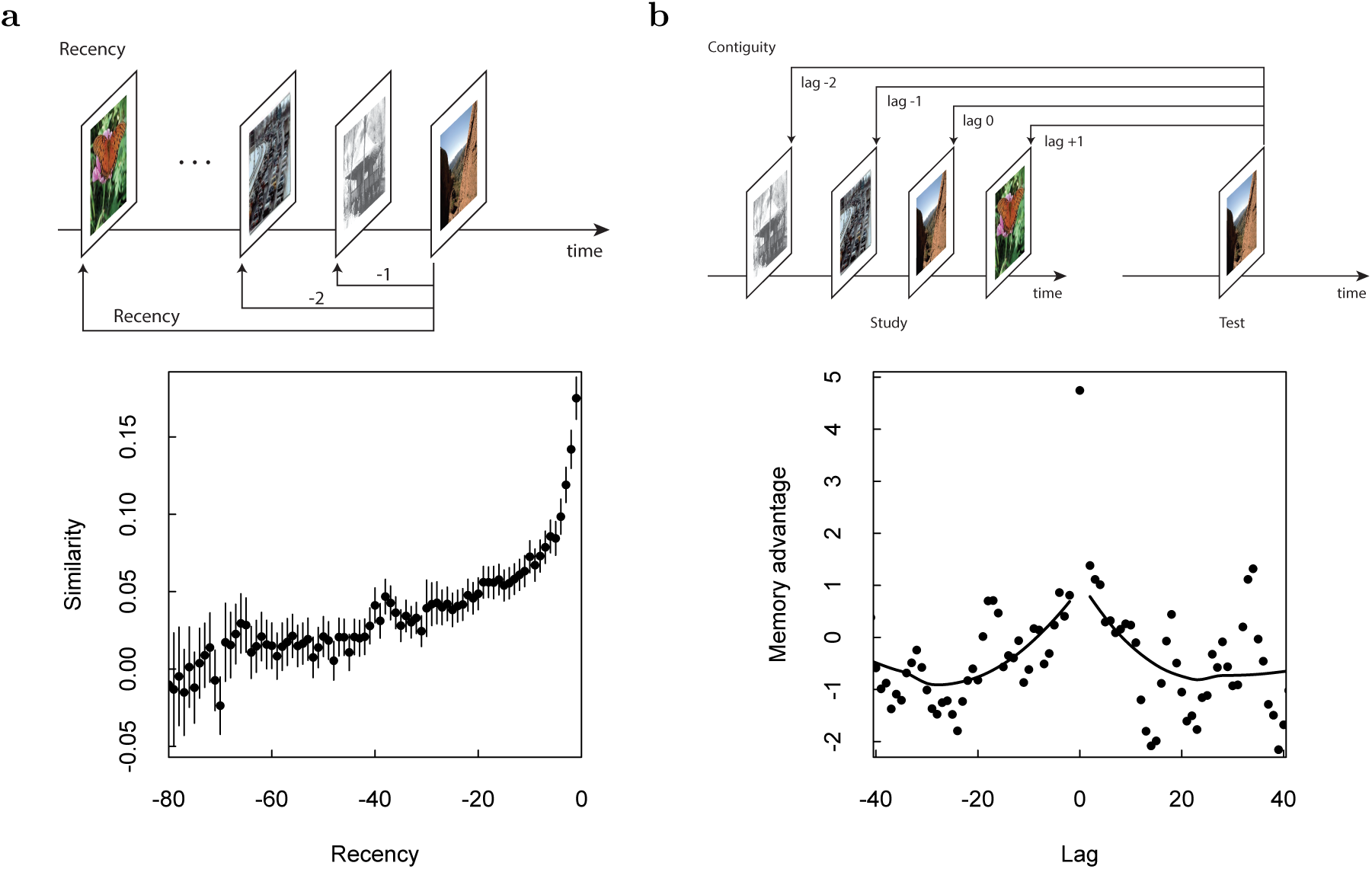
Neural jump-back in time. **a.** Neural recency effect. Top: Schematic describing the definition of recency. For each presentation of a stimulus, a population vector was computed for the 2 s following presentation of the stimulus. This vector was then compared to the population vector from all preceding stimulus presentations and the similarity was aggregated as a function of the recency between the comparisons. Bottom: The population vector shows a recency effect, changing (conservatively) to at least recency −30 during study, corresponding to about two minutes. Smoothed curves are from a LOESS regression. **b.** Neural contiguity effect showing a jump-backin-time. Top: Schematic of the lag variable. For a test probe, similarity of the population vector after the test probe is compared to the population vectors of each study event. The similarity is aggregated as a function of lag, the difference between the original presentation of the probe stimulus and the other list stimulus; the lag to the repeated stimulus is zero. Bottom: To isolate the effect due to episodic memory, we took the difference between the similarity for pictures receiving a highest-confidence response and pictures that were not well-remembered (see methods for details). This “memory advantage” is in units of a paired t-statistic. For clarity, a sliding binning procedure was used to plot the results for lags other than zero. Critically, the memory advantage is peaked around zero, falling off gradually in both the forward and backward directions, indicating a neural jump-back-in-time associated with successful episodic memory retrieval.

We next tested whether this neural recency effect was also visible for specific subsets of visually selective (VS) units (Rutishauser et al., 2015). As Rutishauser et al. (2015) described previously, VS units respond shortly after the onset of a stimulus conditional on the visual category of a stimulus. For example, a subset of “animal selective” VS units change their firing rate only when the image contains an animal. 213 (out of 1286) recorded units qualified as VS units (see methods) and the analysis that follows is restricted to these units. If VS units only carry information about the stimulus that was just presented, their activity should not vary gradually and would thus not show a temporal context effect. Contrary to this prediction, we found that VS units also exhibited a robust neural similarity effect similar to that observed to all recorded units (Fig. 8a). This was true both during study, .0013 ±.0001, *F*_(1,78)_ = 104.6, *p* < .001, as well as test, .0009 ±.0001, *F*_(1,78)_ = 84.64, *p* < .001. Consequently, the response of VS units is modulated by temporal context in addition to visual input. This suggests that feedforward visual input is modulated by temporal context, a critical prediction of the temporal context model.

### Episodic memory was associated with the recovery of temporal context

We next computed the similarity in neural response between pairs of test and study items (see Figure 6b, top, for an illustration). The contextual reinstatement model predicts that the neural response to a recollected old probe that was originally presented at position *i* will be similar to the neural response to study events that were presented shortly before or after position *i*. The variable lag describes the difference between the serial position of the original presentation of an old probe and a study item; lag zero corresponds to the comparison between an old probe at test and its original presentation during study (when it was new). We would expect the neural pattern similarity at lag zero to be large to the extent the response is determined by visual input, which is similar for study and test of the same stimulus.

### Raw contiguity analyses

We first computed the neural similarity as a function of lag separately for recollected probes and unrecollected probes—operationalized as probes that did and did not receive a highest confidence old response (Figure 7).^4^

**Figure 7.**
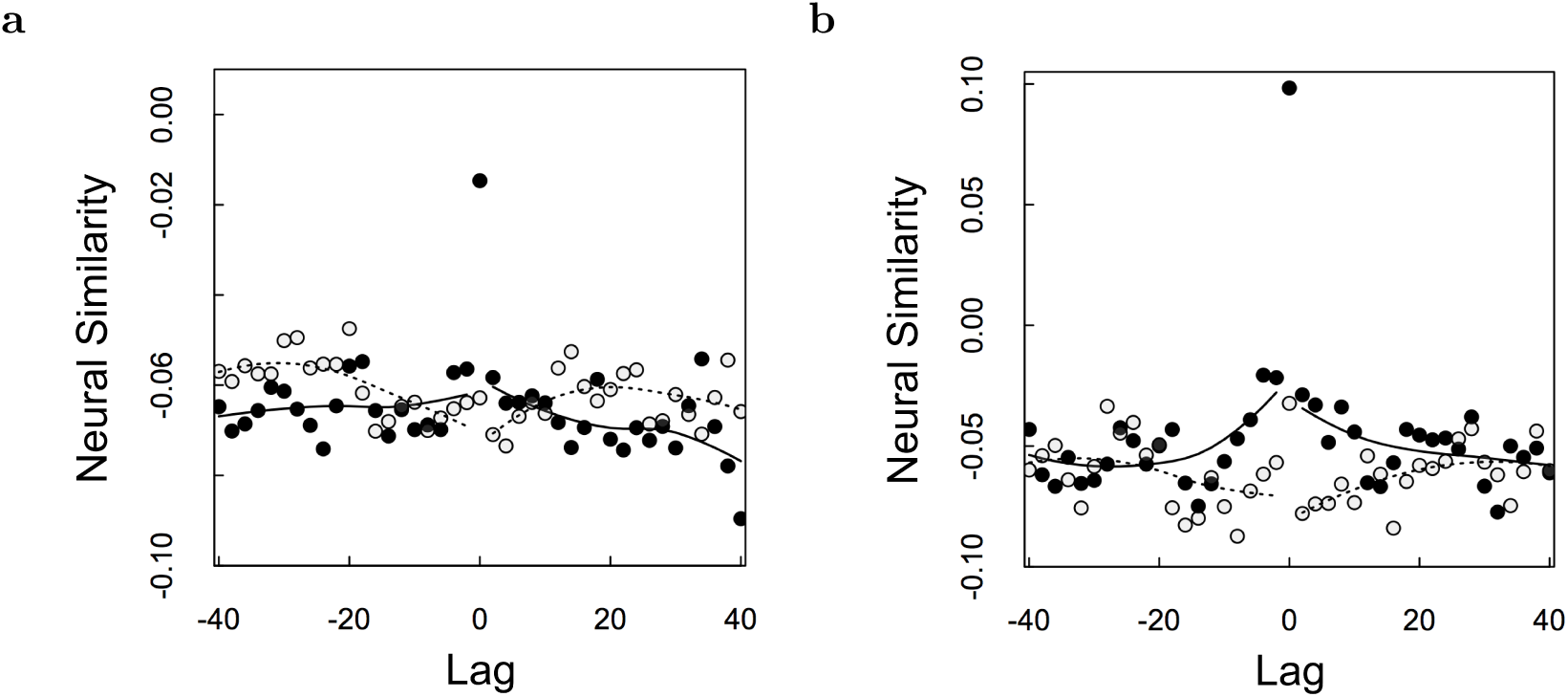
Neural similarity as a function of lag for old probes that received a highest-confidence yes response (filled circles) and old probes that did not receive a highest-confidence yes response (grey open circles). Statistical analyses confirm that there was a contiguity effect (inverted-V centered around zero) for remembered probes but an anti-contiguity effect (V-shaped centered around zero) for unremembered probes. All data points except lag zero were binned. A LOESS curve was fitted for each data set. **a.** All units. **b.** Analysis restricted to units categorized as visually-selective.

The similarity at lag zero was much higher than other lags for highest-confidence responses for both all units taken together and for VS units taken alone. For all units (Fig. 7a), the similarity value at lag zero for probes that received a highest-confidence response (filled circles) was greater than for other lags (sign test, *p* < .001). Similar results were found for VS units taken alone (Fig. 7b). For old probes that did not receive a highest-confidence response (open circles), there was no discernible advantage for lag zero for all units taken together. For VS units there was a reliable advantage for lag zero over other lags (*p* < .001 by a sign test) although the numerical value was much smaller than for probes that received a highest-confidence response. This effect at lag zero is unsurprising and would be expected to hold for any neural process that responds to the features of the stimuli.

In addition similarity tended to decrease as a function of |lag|for old probes that received a highest confidence response. That is, for old probes that received a highest confidence response, the population vector at test was more similar to the population vector for study events close in time—in both the forward and backward direction—to the time at which that probe stimulus was originally studied. In contrast, there was a tendency for similarity to *increase* as a function of |lag*|* for probes that were not recollected. That is, for probes that did not receive a highest confidence response, the population vector at test was *less* similar to the population vectors for items studied close in time to the probe stimulus than for items studied further away. This suggests that the degree of contextual reinstatement predicted the success or failure of episodic retrieval. Both of these findings also held (see Fig. 7b) for VS units considered alone.

We quantified this by performing an ANOVA with |lag*|* as regressor (excluding lag zero) and direction (backward or forward) as a categorical variable separately for old probes that did and did not attract a highest-confidence old recognition judgment. For simplicity we will refer to these as recollected and unrecollected probes. We did this ANOVA for both all units (Fig. 7a) and restricting our attention to VS units (Fig. 7b). Both set of analyses led to similar conclusions. Considering all units, the neural similarity for recollected probes showed a significant effect of |lag*| F*_(1,56)_ = 8.09, *p* < .01 and no effect of direction *F*_(1,56)_ = 2.15, nor an interaction *F*_(1.56)_ = 1.40. For unrecollected probes the effect of |lag*|* was again significant *F*_(1,56)_ = 9.59, *p* < .005, and there was neither an effect of direction *F*_(1,56)_ = 1.35, nor an interaction *F*_(1.56)_ = 1.79. However, the effect of |lag*|* was in different directions for recollected and unrecollected probes. For recollected probes, the effect of |lag*|* was positive in both the forward and backward directions (like an inverted-V); for unrecollected probes, the effect of |lag*|* was negative in both the forward and backward directions (a V-shaped curve around zero). For recollected old probes, the effect of lag on neural similarity in the forward direction reached significance, (-.46 ± .15) *×* 10^−3^, *p* < .005 and the contiguity effect in the backward direction did not reach significance (.19 ±.18) *×* 10^−3^, *p* > .2. For unrecollected probes there was a reliable negative effect of lag in the backward direction, (-.52 ±.17) *×* 10^−3^, *p* < .005. There was a similar trend in the forward direction, although the trend did not reach significance (.21 ± .16) *×* 10^−3^.

Furthermore, there was an interaction such that recollected probes showed a different dependency on |lag*|* than unrecollected probes. Using |lag*|* as the regressor and recollection as a categorical variables an ANOVA showed a significant interaction term *F*_(1,116)_ = 17.27, *p* < .001.

### Subdividing probes that did not receive a highest-confidence response

The preceding analysis compares the neural contiugity effect to old probes that received a highest-confidence old response to the neural contiguity effect to all other old probes. In that analysis, old probes that received a lower-confidence old response (4-5 on the six-point scale) are collapsed with misses, old probes that received a new response (1-3 on the six point scale). To determine if there were reliable differences in the neural contiguity effect for probes that received a 4-5 response we conducted further analyses comparing these three categories of responses to one another.

There was no evidence of an effect of |lag*|* for old probes that received a response of confidence 4-5, *F*_(1,58)_ = 1.615. The contiguity effect for probes that received a 4-5 response was reliably different from the neural contiguity effect for probes that received a highest-confidence response; an ANOVA with |lag*|* as a regressor and confidence level (4-5 *vs* 6) as a categorical variable showed a significant interaction term between confidence level and |lag|, *F*_(1,116)_ = 5.65 *p <* 0.05. Further, we investigated if there is a significant difference between the neural contiguity effect caused by familiar old items (4-5 on the six point scale) and to probes rated as new (1-3 on the six-point scale). An ANOVA similar to the above did not show a significant interaction between the two groups *F*_(1,116)_ = 0.1646. In light of these results, our subsequent analyses only compared old probes that received highest-confidence responses (6 on a six point scale) to all other old probes (1-5 on a six point scale).

### Episodic memory was associated with a neural jump-back-in-time

To isolate the contribution to neural pattern similarity attributable to episodic memory, we calculated the difference between the neural pattern similarity as a function of lag for probes that received a highest-confidence response and those that did not (see methods for details). In the following, we refer to this difference as ‘memory advantage,’ which is in units of a *t*-statistic comparing recollected to unrecollected probes. By examining the memory advantage as a function of lag, we can simply assess whether episodic memory retrieval is associated with a neural contiguity effect as predicted by retrieved context models.

The memory advantage index showed a robust contiguity effect (Figure 6b). This was also true for VS units considered alone (Figure 8b). The memory advantage at lag zero was significant, *t* = 4.75, *p* < .001. The effect at lag zero, however, does not indicate reinstatement of temporal context. This is because similar visual features were present during study of stimulus *i* and test of stimulus *i*. To test for a neural jump-back-in-time, we asked whether the memory advantage changed systematically as a function of lag. A jumpback-in-time requires that the repeated image presentation triggers a retrieval of previous context, i.e. the reinstatement of the neural ensemble activity present before the first encounter with the probe stimulus. This would be expected to manifest in a decrease in the neural memory advantage as a function of lag in both the forward direction (lag increasing from zero) and in the backward direction (lag decreasing from zero).

**Figure 8.**
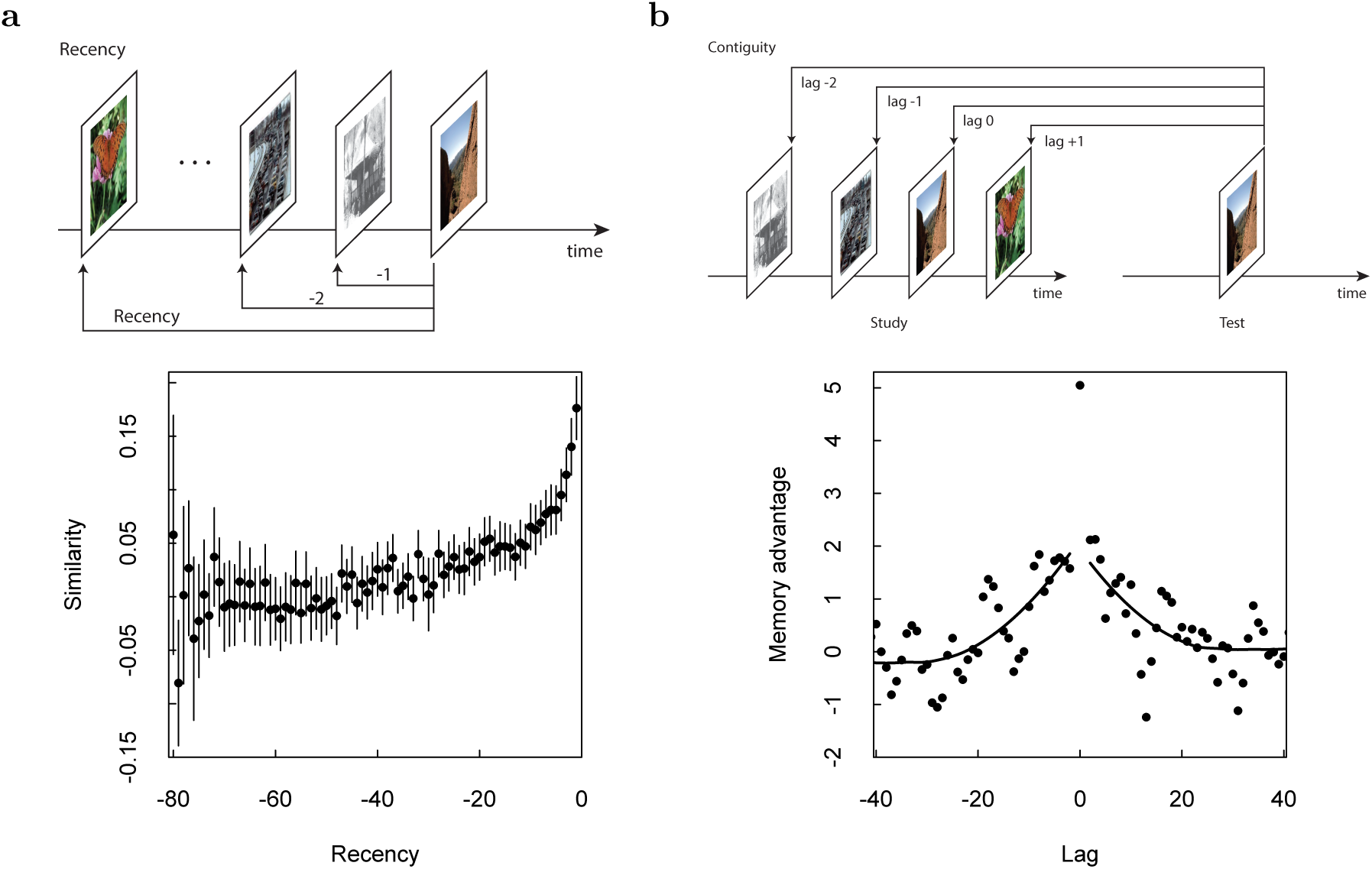
Visual-category sensitive units showed neural recency and contiguity effects. Format is as in Fig. 6b, but with analyses restricted only to units that differentiated the category of the currently-presented image during study.

#### Parametric analyses

To evaluate this prediction we performed an ANOVA on memory advantage with |lag*|* as regressor (excluding lag zero) and direction (backward or forward) as a categorical variable. There was a significant effect of |lag|, *F*_(1,56)_ = 16.4, *p* < .001 but no effect of direction, *F*_(1,56)_ = .003 nor an interaction *F*_(1,56)_ = .01. The effect of |lag*|* means that the similarity of the population vector after recovery to the population vectors close together in time to the original presentation of the probe stimulus predicted whether the probe triggered an episodic memory (attracted a highest confidence old judgement). This is as predicted by the hypothesis that episodic memory is accompanied by recovery of a gradually-changing state of temporal context in the human brain.

A decrease in the memory advantage extending to lags near zero but restricted to the forward direction (positive values of lag) could correspond to persistence of stimulus-specific features in memory such as in a short-term memory buffer. In contrast a jump-back-in-time would cause reconstruction of the pattern of activation *prior* to initial presentation of the probe. Thus, a jump-back-in-time would manifest as a advantage for lags near zero in *both* directions. To quantify the effect, we performed linear regressions of memory advantage onto lag separately for each direction. We found reliable regression coefficients for both the forward (lags 1 to 30) and backward (lags −30 to −1) direction (forward: −.06 ± .02, *F*_(1,28)_ = 7.69, *p* < .01; backward: .07 ± .02, *F*_(1,28)_ = 8.694, *p* < .01). Thus, we found an effect of lag on the memory index separately in both the forward and backward directions as predicted by a neural jump-back-in-time.

#### Permutation analysis

In order to determine if these results were simply due to violation of one or more assumptions of the parametric tests, we also performed permutation tests to evaluate the probability of obtaining these results by chance. The results of the permutation analysis was consistent with the conclusions from the parametric statistics. The observed regression coefficient for |lag*|* (.06) was more extreme than the regression coefficient for 997/1000 permutations. The permuted regression coefficients were approximately normal −.004 ± .02 (mean *±* SD). In addition to the regression of |lag|, we also computed the regression coefficients for the forward and backward regressions on the permuted data. The observed value for the forward and backward regression coefficients (−.06 and .07 respectively) were more extreme than 970/1000 and 974/1000 of the values from the permuted data. Again the permuted statistics were approximately normally distributed, −.001 ± .03 and .008 ± .03 respectively.

### Visually-selective units showed a jump-back-in-time

We found similar evidence for a neural jump-back-in-time when considering only VS units (Figure 8b). The memory advantage at lag zero was significant, *t* = 5.05, *p* < .001. An ANOVA with |lag*|* as regressor (excluding lag zero) and direction (backward or forward) as a categorical variable showed a significant effect of |lag|, *F*_(1,56)_ = 18.9, *p* < .001 but no effect of direction, *F*_(1,56)_ = .07 nor an interaction, *F*_(1,57)_ = .91.

Also, considering the forward and backward directions separately, we found a regression coefficient of −.06 ±.03, *F*_(1,28)_ = 5.4, *p* < .03 for the forward direction (lag 1 to lag 30) and .09 ± .02, *F*_(1,28)_ = 15.03, *p* < .001 for the backward direction (lag −30 to lag −1). Thus, a signal compatible with contextual reinstatement was visible even when only considering VS units that were sensitive to the category of the visual stimulus presented during study.

The conclusions of these parametric analyses of contiguity restricted to the VS units were also supported by the results of the permutation analysis. The true regression coefficient for |lag|, −.08, was more extreme than the coefficients from all 1000 permutations. The regression coefficents for forward and backward lag (-.06 and .09 respectively) were greater than the values for 966/1000 and 993/1000 permutations.

Although the effect of contiguity was significant for the VS units taken in isolation, there was not a reliable difference between the contiguity effect in the memory advantage index for VS units and the contiguity effect in the memory advantage index for non-VS units. An ANOVA with |lag*|* as a regressor and group (VS/not-VS) as a categorical variable showed main effects of |lag|, *F*_(1,116)_ = 25.27, *p <* 0.001 and group *F*_(1,116)_ = 39.96, *p <* 0.001 but no interaction of |lag*|* and group *F*_(1,116)_ = 2.72.

### Contiguity signal separated by brain region

Separating the analysis according to brain regions, we found that units in the amygdala and hippocampus displayed a contiguity effect taken in isolation. In addition, there was not evidence that the contiguity effect differed between the two brain regions, supporting its existence independently in both areas.

An ANOVA of the memory advantage in the amygdala showed a reliable effect of |lag|, *F*_(1,56)_ = 17.14, *p <* 0.001, without an effect of direction, *F*_(1,56)_ = 0.558, nor an interaction between |lag*|* and direction, *F*_(1,56)_ = 0.0009. The main effect of |lag*|* was substantiated by a permutation analysis; the observed regression coefficient was larger than 1000/1000 values from permutations of the trials. Furthermore, the effect of decreasing similarity is evident in both directions, with the regression analysis in the forward direction yielding a significant coefficient in both the forward, −0.07 *±* 0.03, *p <* 0.05 and backward, lag = 0.07 *±* 0.02, *p <* 0.001, directions. These parametric results within the amygdala were supported by permutation analyses; the observed values were more extreme than 980/1000 and 986/1000 values from shuffled data.

Considering the hippocampus in isolation, we also observed a contiguity effect on the memory index. There was a main effect of |lag|, *F*_(1,58)_ = 12.13, a weakly significant effect of direction *F*_(1,58)_ = 4.93, *p* < .05, and no interaction between direction and lag *F*_(1,58)_ = 0.4217. The observed regression coefficient for |lag*|* was greater than 978/1000 values from permuted data. Although the forward direction yielded a reliable regression coefficient, lag = −0.06 *±* 0.02, *p <* 0.01, the regression coefficient in the backward direction did not reach significance 0.04 *±* 0.02, *p >* 0.09. The observed regression coefficient in the forward direction exceeded 942/1000 shuffled values. The observed regression coefficient in the backward direction exceeded 887/1000 shuffled values.

Most importantly, we did not find strong evidence that the effect of contiguity on the memory index was different across regions. An ANOVA with |lag*|* as regressor and brain region (amygdala *vs* hippocampus) as categorical variable, showed a significance of brain region *F*_(1,116)_ = 7.1163 *p <* 0.01, but not a significant interaction *F*_(1,116)_ = 0.7259. A permutation analysis showed that in 547/1000 cases the permuted data showed an interaction term this large.

These results are consistent with the hypothesis that the amygdala and the hippocampus have the same contiguity effect but that it is more difficult to measure in the hippocampus than in the amygdala. Indeed the recordings yielded about 1.6 times as many units from the amygdala than in the hippocampus. Of the 1286 units used for the analyses, 800 were located in the amygdala and 486 in the hippocampus.

### Additional evidence for a jump back in time

If some old probes caused a jump back in time, we would expect this to result in greater pattern similarity between pairs of test vectors and pairs of study vectors. If two test probes recover information from the temporal context during presentation of those items, then this would be expected to result in additional similarity between these two test events if those two test probes were close together in the list. Consistent with this hypothesis, although both study and test lists showed a reliable recency effect, for the test list the similarity stablized at a higher baseline value. At recency less than about −20, the test similarity was reliably higher (Fig. 9) at almost all values of recency.

As a simple quantitative measure the similarity for test is greater than the similarity for study at 13/20 of the points for recency −1 through −20. In contrast, for recency −21 through −80 the similarity is greater for 55*/*60 of the values. These two proportions differ from one another, *χ*^2^(1) = 6.41, *p* < .05.

**Figure 9.**
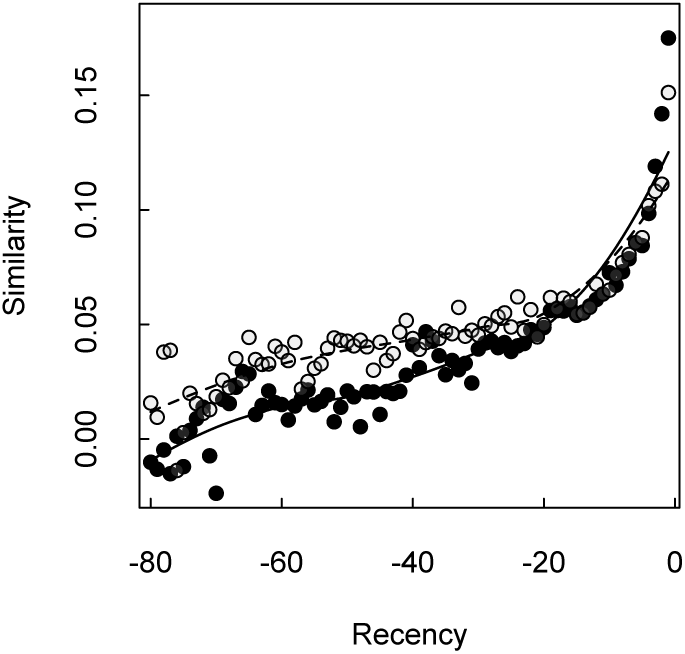
Enhanced neural similarities during test. Neural recency effects for study (filled circles) and test (open circles). The neural similarity was higher between test events than between study events over a wide range of values of recency. This advantage consistent with the predictions of a retrieved temporal context model. Smoothed curves are from a LOESS regression.

## Discussion

We found that the population activity of human MTL units changed gradually over minutes (Fig. 6a). This replicates, in humans, prior evidence for a gradually-changing temporal context signal found previously in the MTL of animals (Naya & Suzuki, 2011; MacDonald et al., 2011; Manns et al., 2007; Hyman et al., 2012; Mankin et al., 2012; Cai et al., 2016; Rashid et al., 2016; Rubin et al., 2015) and humans (Howard et al., 2012; Yaffe et al., 2014; Manning et al., 2011; Hsieh, Gruber, Jenkins, & Ranganath, 2014; Hsieh & Ranganath, 2015). Crucially, visually selective category cells also showed this gradual change (Fig. 8a). This is important because it suggests that the population vector does not merely change gradually over time but also carries information about the identity of the stimuli presented.

The critical new insight that this paper contributes is a first demonstration that the retrieval of human episodic memory is associated with the recovery of a gradually-changing state of temporal context—a neural jump back in time. This analysis measures the similarity between a population vector caused by an old probe and population vectors during study of the neighbors of the original presentation of the probe stimulus. The difference between the population similarity calculated for probes that triggered an episodic memory (operationalized as a highest-confidence old response) and the population similarity calculated for probes that did not trigger an episodic memory was greater for the neighbors of the original presentation and fell off reliably with distance from the original presentation of the probe stimulus in both directions (Fig. 6b). Notably, a robust contiguity effect associated with memory was observed when considering population vectors constructed from only the visual selective units (Fig. 8b).

We did not merely observe contiguity effect for recollected probes, but also an anti-contiguity effect for old probes that were not recollected (Fig. 7). Methodologically, this means that had we averaged over all old probes, it would have been very difficult to observe a neural contiguity effect. Had we observed that the probes that did not evoke a highest-confidence response resulted in a contiguity effect that was weaker, this would suggest a continuity between recollection and familiarity. Because there was not such an effect, and indeed a tendency towards an anti-contiguity effect it suggests that in this study successful retrieval of preceding temporal context is only observed for probes that received a highest-confidence response. Put another way, if the degree of reinstatement of the neural population causes a high-confidence response, we would expect that high-confidence probes would correspond to a high degree of reinstatement *and* probes that received a lower confidence response correspond to a low degree of reinstatement. This latter property predicts an anti-contiguity effect.

In this study, we observed that the brain state in the moments after retrieval of an episodic memory resembled the gradually-changing temporal context at the time that memory was encoded. However, this does not imply that this recovered context persists long after the recollection of that probe. If context retrieved by a probe persisted long after the presentation of the probe, one would expect the rate of contextual drift during the test list to be very different than the rate of drift during study. While there is some difference (Figure 9), the discrepancy is modest compared to what one would expect if retrieved context persisted long after the presentation of a probe. One possibility is that retrieved context only becomes available for a short time after presentation of the probe and then dissipates. This is analogous with “awake replay” events in the rodent hippocampus (Carr, Jadhav, & Frank, 2011; Pfeiffer & Foster, 2015), in which hippocampal place cells briefly fire as if the animal is in a remote location during sharp-wave-ripple events. Perhaps the neural jump-back-in-time is a transient discontinuity in the stream of temporal context much like the transient discontinuity in the representation of position.

### Methodological advantages of the present study compared to previous attempts to measure a neural jump-back-in-time

This study avoids methodological pitfalls of previous papers that addressed whether human episodic memory is associated with a jump back in time. A previous study with human single units (Howard et al., 2012) used continuous recognition, in which probes are intermixed with study items in a continuous stream of experience. In that study there was a robust behavioral recency effect. Because recency is confounded with lag in the backward direction, it was necessary to statistically decouple recency from contiguity in that study. Here, the 30 minute delay between study and test and the absence of a behavioral recency effect eliminated any confound due to recency. Note also that we would expect a neural recency effect to be present for old probes that were not recollected as well as old probes that were recollected. Because the neural contiguity effect was observed in the difference between these suggests it is not due to a confound between recency and contiguity.

Another prior study used autocorrelated features from ECoG in a free recall study (Manning et al., 2011). They found that the features during recall of the word studied at serial position *i* in the list resembled the features during study of nearby list items. However, because free recall is extended in time and exhibits a robust behavioral contiguity effect (Kahana, 1996; Sederberg, Miller, Howard, & Kahana, 2010) that finding does not establish that recall of word *i* is associated with *recovery* of a gradually changing temporal context. Because of the behavioral contiguity effect, the recall of word *i* is likely to have been preceded by neighbors of word *i*, so that the neural contiguity effect could have been due to the persistence of item representations from previous recalls. Similar concerns apply to a human single unit study that argued for recovery of spatial context during a free recall task (Miller et al., 2013). Another ECoG study used cued recall to establish that successful recovery of a word was associated with recovery of temporally-varying features from the list (Yaffe et al., 2014). Although this study was able to establish a correlation between successful memory and a contiguity effect, the analyses included lag zero in the measurement of the neural contiguity effect in the backward direction. Because the neural contiguity effect in the backward direction is a distinctive signature of a jump back in time, whereas similarity at lag zero may be attributed to repeated items, the analyses reported in that paper did not clearly establish a neural jump back in time.

### Recent studies showing the importance of temporal context in human memory

This study adds to a growing body of work from human cognitive neuroscience that suggests that a gradually-changing state of temporal context affects memory in a range of tasks. A free recall study using fMRI showed that the content of lingering item representations during study predicted free recall transitions during retrieval (Chan, Applegate, Morton, Polyn, & Norman, in press). A recent fMRI study showed that the amount of drift in the right entorhinal cortex between two events in a radio program predicted participants’ judgment of the duration between the two events (Lositsky et al., 2016). Similarly, when participants rate the relative recency of two probes, hippocampal pattern similarity predicted the order judgment (DuBrow & Davachi, 2014, 2016). Moreover, in that same study, successful judgments were associated with reinstatement of stimuli that intervened between the two probes. A recent study with patients with MTL damage showed that patients were impaired at their ability to perform temporal ordering, as if an intact MTL was required for recovering temporal context (Dede, Frascino, Wixted, & Squire, 2016).

Finally, a pair of recent studies suggest that the recovery of temporal context we observed in the laboratory could also reflect a mechanism for memory in more natural settings. In natural settings, the visual features the participant experiences are autocorrelated in both time and space, unlike the randomly-assembled list of visual stimuli experienced in a fixed location used in the present study. A recent study of natural memory automatically recorded pictures as participants went about their daily lives for several weeks (Nielson, Smith, Sreekumar, Dennis, & Sederberg, 2015). After a delay, participants were brought into the scanner and shown images from their own lives. The pattern similarity between pairs of images that were well-remembered was computed. The pattern similarity in the anterior hippocampus predicted the distance in both time and space between pairs of remembered images, on the scale of hours to weeks for time and tens of meters to kilometers for space. Another recent study, adding to work in virtual reality environments (Chadwick, Hassabis, Weiskopf, & Maguire, 2010; Copara et al., 2014) observed similar results for episodic memory in a well-controlled virtual environment in which spatial and temporal proximity could be deconfounded (Deuker, Bellmund, Schröder, & Doeller, 2016). In light of this growing body of evidence and modeling work suggesting a deep connection between temporal context and spatial context (Howard et al., 2014; Howard & Eichenbaum, 2015), the present study suggests that recovery of a gradually-changing state of spatiotemporal context is an essential aspect of human episodic memory that depends crucially on the function of the MTL.

### Implications for theory of hippocampus and episodic memory

We found that the activity of populations in the human MTL changed gradually over at least a minute, adding to a large and growing body of evidence that neural states change gradually in the MTL (e.g., Manns et al., 2007; Mankin et al., 2012; Hyman et al., 2012; Cai et al., 2016; Rashid et al., 2016, see Howard, 2017 for a review). The present results suggest that these gradually-changing states are recovered during retrieval of an episodic memory. This provides a challenge for traditional models of memory retrieval that rely on autoassociative pattern completion (e.g., Hopfield, 1982). Attractor models of content-addressable memory are notoriously sensitive to overlap in the stored patterns and the capacity of attractor networks plummets if the stored patterns are correlated with one another (e.g., Amit, Gutfreund, & Sompolinsky, 1985). Traditionally, the way this inherent tension has been addressed is to assume that the pattern completion stage performed by the attractor network is preceded by a pattern separation stage that decorrelates the patterns before they are stored (Marr, 1971; Levy, 1989; McClelland, McNaughton, & O’Reilly, 1995). This hypothesis has been extremely influential and has inspired a wealth of empirical work, especially cognitive neuroimaging work (e.g., Bakker, Kirwan, Miller, & Stark, 2008). If pattern similarity was a big problem for the computational mechanism used to recover a memory one might have expected the brain to avoid pattern similarity at all costs. However, even in a randomly-assembled list of pictures, the brain induces robust overlap, introducing pattern similarity to the neural states extending at least a minute. One possibility is that the CA3 field of the hippocampus is decorrelated, suggesting that populations in CA3 would not change gradually. Although we are unable to identify the subfields of the hippocampus in this study, animal work has provided mixed results addressing this point. Mankin et al. (2012) observed less temporal drift in CA3 than in CA1 over long periods of time during open field foraging tasks. However, Salz et al. (2016) observed robust sequences of time cells in the CA3 that were indistinguishable from those observed in CA1 in the same experiment.

It is also possible that pattern separation followed by pattern completion is not necessary because the hippocampus does not rely on an autoassociative content-addressable memory (Teyler & DiScenna, 1985, 1986). Rather than relying on content-addressable pattern completion, time and space could function like pointers for an address-addressable memory (Howard, in press). In this view, the hippocampus contains a map of indexes to the content that can be found at different temporal and spatial addresses. Singh, Oliva, and Howard (2017) measured the amount of time to access memory for a picture as a function of how far in the past it was experienced in a continuous recognition experiment. They found that the time to access a memory went up linearly with the logarithm of the recency of the probe stimulus. These results are as one would expect if memory accessed depended on scanning across a logarithmically compressed timeline of the past (Howard et al., 2015).

## Footnotes

1 The complete dataset includes sixty-five sessions. It includes sessions that became available after the analyses in the current paper were begun.

2 Subsequent analyses subdivided old probes that received a lower-confidence old response (4-5 on the six-point scale) from old probes that received a new response (1-3 on the six-point scale).

3 In two sessions, the participant only used the two extreme response keys for the new probes, making it impossible to measure a slope.

4 The negative values for the contiguity analysis do not imply that the vectors are anti-correlated. To see why, start with a skewed random variable and take the z-score of this variable. The expected value of the product of independent draws from this z-scored random variable is negative. Negative values of the contiguity analysis are a consequence of the skew of the distribution of firing rates.

## References

Amit, D. J., Gutfreund, H., & Sompolinsky, H. (1985). Storing infinite numbers of patterns in a spin-glass model of neural networks. Physical Review Letters, 55 (14), 1530–1533.

Bakker, A., Kirwan, C. B., Miller, M., & Stark, C. E. (2008). Pattern separation in the human hippocampal ca3 and dentate gyrus. Science, 319 (5870), 1640–2.

Cai, D. J., Aharoni, D., Shuman, T., Shobe, J., Biane, J., Song, W.,… Silva, A. (2016). A shared neural ensemble links distinct contextual memories encoded close in time. Nature, 534 (7605), 115–118.

Carr, M. F., Jadhav, S. P., & Frank, L. M. (2011). Hippocampal replay in the awake state: a potential substrate for memory consolidation and retrieval. Nature Neuroscience, 14 (2), 147– 153.

Chadwick, M. J., Hassabis, D., Weiskopf, N., & Maguire, E. A. (2010). Decoding individual episodic memory traces in the human hippocampus. Current Biology, 20 (6), 544–547.

Chan, S. C. Y., Applegate, M. C., Morton, N. W., Polyn, S. M., & Norman, K. A. (in press). Lingering representations of stimuli influence recall organization. Neuropsychologia.

Copara, M. S., Hassan, A. S., Kyle, C. T., Libby, L. A., Ranganath, C., & Ekstrom, A. D. (2014). Complementary roles of human hippocampal subregions during retrieval of spatiotemporal context. Journal of Neuroscience, 34 (20), 6834–6842.

Dede, A. J., Frascino, J. C., Wixted, J. T., & Squire, L. R. (2016). Learning and remembering real-world events after medial temporal lobe damage. Proceedings of the National Academy of Sciences, 113 (47), 13480–13485.

Deuker, L., Bellmund, J. L., Schr¨oder, T. N., & Doeller, C. F. (2016). An event map of memory space in the hippocampus. eLife, 5, e16534.

Diana, R. A., Yonelinas, A. P., & Ranganath, C. (2007). Imaging recollection and familiarity in the medial temporal lobe: a three-component model. Trends in Cognitive Science, 11 (9), 379–86.

DuBrow, S., & Davachi, L. (2014, Oct). Temporal memory is shaped by encoding stability and intervening item reactivation. Journal of Neuroscience, 34 (42), 13998–4005. doi:10.1523/JNEUROSCI.2535-14.2014

DuBrow, S., & Davachi, L. (2016). Temporal binding within and across events. Neurobiology of learning and memory, 134, 107–114.

Eichenbaum, H., Yonelinas, A., & Ranganath, C. (2007). The medial temporal lobe and recognition memory. Annual Review of Neuroscience, 30, 123–152.

Faraut, M., Carson, A., Sullivan, S., Tudoscicuc, O., Ross, I., Reed, C.,… Rutishauser, U. (2018a). Data from: Dataset of human medial temporal lobe single neuron activity during declarative memory encoding and recognition. https://doi.org/10.5061/dryad.46st5.

Faraut, M., Carson, A., Sullivan, S., Tudoscicuc, O., Ross, I., Reed, C.,… Rutishauser, U. (2018b). Dataset of human medial temporal lobe single neuron activity during declarative memory encoding and recognition. Scientific Data, 5, 180010 EP.

Fortin, N. J., Wright, S. P., & Eichenbaum, H. (2004). Recollection-like memory retrieval in rats is dependent on the hippocampus. Nature, 431 (7005), 188–91.

Hassabis, D., Kumaran, D., Vann, S. D., & Maguire, E. A. (2007). Patients with hippocampal amnesia cannot imagine new experiences. Proceedings of the National Academy of Sciences USA, 104 (5), 1726–31. doi:10.1073/pnas.0610561104

Hautus, M. J., Macmillan, N. A., & Rotello, C. M. (2008). Toward a complete decision model of item and source recognition. Journal of Experimental Psychology: Learning, Memory, and Cognition, 15, 889–905.

Hopfield, J. J. (1982). Neural networks and physical systems with emergent collective computational abilities. Proceedings of the National Academy of Science, USA, 84, 8429–8433.

Howard, M. W. (2017). Temporal and spatial context in the mind and brain. Current Opinion in Behavioral Sciences, 17 (14-19).

Howard, M. W. (in press). Memory as perception of the past: Compressed time in mind and brain. Trends in Cognitive Sciences.

Howard, M. W., & Eichenbaum, H. (2015). Time and space in the hippocampus. Brain Research, 1621, 345–354.

Howard, M. W., & Kahana, M. J. (2002). A distributed representation of temporal context. Journal of Mathematical Psychology, 46 (3), 269–299.

Howard, M. W., MacDonald, C. J., Tiganj, Z., Shankar, K. H., Du, Q., Hasselmo, M. E., & Eichenbaum, H. (2014). A unified mathematical framework for coding time, space, and sequences in the hippocampal region. Journal of Neuroscience, 34 (13), 4692–707. doi:10.1523/JNEUROSCI.5808-12.2014

Howard, M. W., Shankar, K. H., Aue, W., & Criss, A. H. (2015). A distributed representation of internal time. Psychological Review, 122 (1), 24–53.

Howard, M. W., Viskontas, I. V., Shankar, K. H., & Fried, I. (2012). Ensembles of human MTL neurons “jump back in time” in response to a repeated stimulus. Hippocampus, 22 (9), 1833– 1847.

Howard, M. W., Youker, T. E., & Venkatadass, V. (2008). The persistence of memory: Contiguity effects across several minutes. Psychonomic Bulletin & Review, 15 (PMC2493616), 58–63.

Hsieh, L.-T., Gruber, M. J., Jenkins, L. J., & Ranganath, C. (2014). Hippocampal activity patterns carry information about objects in temporal context. Neuron, 81 (5), 1165–1178.

Hsieh, L.-T., & Ranganath, C. (2015). Cortical and subcortical contributions to sequence retrieval: Schematic coding of temporal context in the neocortical recollection network. NeuroImage, 121, 78–90.

Hyman, J. M., Ma, L., Balaguer-Ballester, E., Durstewitz, D., & Seamans, J. K. (2012). Contextual encoding by ensembles of medial prefrontal cortex neurons. Proceedings of the National Academy of Sciences USA, 109, 5086–91. doi:10.1073/pnas.1114415109

Kahana, M. J. (1996). Associative retrieval processes in free recall. Memory & Cognition, 24, 103–109.

Kahana, M. J. (2012). Foundations of human memory. OUP USA.

Levy, W. B. (1989). A computational approach to hippocampal function. In R. D. Hawkins & G. H. Bower (Eds.), Computational models of learning in simple neural systems (p. 243–305). New York: Academic Press.

Lositsky, O., Chen, J., Toker, D., Honey, C. J., Shvartsman, M., Poppenk, J. L.,… Norman, K. A. (2016, nov). Neural pattern change during encoding of a narrative predicts retrospective duration estimates. eLife, 5, e16070. doi:10.7554/eLife.16070

MacDonald, C. J., Lepage, K. Q., Eden, U. T., & Eichenbaum, H. (2011). Hippocampal “time cells” bridge the gap in memory for discontiguous events. Neuron, 71 (4), 737–749.

Mankin, E. A., Sparks, F. T., Slayyeh, B., Sutherland, R. J., Leutgeb, S., & Leutgeb, J. K. (2012). Neuronal code for extended time in the hippocampus. Proceedings of the National Academy of Sciences, 109, 19462–7. doi:10.1073/pnas.1214107109

Manning, J. R., Polyn, S. M., Litt, B., Baltuch, G., & Kahana, M. J. (2011). Oscillatory patterns in temporal lobe reveal context reinstatement during memory search. Proceedings of the National Academy of Science, USA, 108 (31), 12893–7.

Manns, J. R., Howard, M. W., & Eichenbaum, H. B. (2007). Gradual changes in hippocampal activity support remembering the order of events. Neuron, 56 (3), 530–540.

Marr, D. (1971). Simple memory: a theory for archicortex. Philosophical Transactions of the Royal Society B, 262 (841), 23–81.

McClelland, J. L., McNaughton, B. L., & O’Reilly, R. C. (1995). Why there are complementary learning systems in the hippocampus and neocortex: insights from the successes and failures of connectionist models of learning and memory. Psychological Review, 102 (3), 419–57.

Miller, J. F., Neufang, M., Solway, A., Brandt, A., Trippel, M., Mader, I.,… Schulze-Bonhage, A. (2013). Neural activity in human hippocampal formation reveals the spatial context of retrieved memories. Science, 342 (6162), 1111–4. doi:10.1126/science.1244056

Naya, Y., & Suzuki, W. (2011). Integrating what and when across the primate medial temporal lobe. Science, 333 (6043), 773–776.

Nielson, D. M., Smith, T. A., Sreekumar, V., Dennis, S., & Sederberg, P. B. (2015). Human hippocampus represents space and time during retrieval of real-world memories. Proceedings of the National Academy of Sciences, 112 (35), 11078–11083.

Onyper, S. V., Zhang, Y. X., & Howard, M. W. (2010). Some-or-none recollection: Evidence from item and source memory. Journal of Experimental Psychology: General, 139 (2), 341–64.

Pfeiffer, B.E., & Foster, D. J. (2015). Autoassociative dynamics in the generation of sequences of hippocampal place cells. Science, 349 (6244), 180–183.

Polyn, S. M., Norman, K. A., & Kahana, M. J. (2009). A context maintenance and retrieval model of organizational processes in free recall. Psychological Review, 116, 129–156.

Rashid, A. J., Yan, C., Mercaldo, V., Hsiang, H.-L. L., Park, S., Cole, C. J.,… others (2016). Competition between engrams influences fear memory formation and recall. Science, 353 (6297), 383–387.

Rubin, A., Geva, N., Sheintuch, L., & Ziv, Y. (2015). Hippocampal ensemble dynamics timestamp events in long-term memory. eLife, 4, e12247.

Rutishauser, U., Aflalo, T., Rosario, E. R., Pouratian, N., & Andersen, R. A. (2017). Single-neuron representation of memory strength and recognition confidence in left human posterior parietal cortex. Neuron.

Rutishauser, U., Mamelak, A. N., & Schuman, E. M. (2006). Single-trial learning of novel stimuli by individual neurons of the human hippocampus-amygdala complex. Neuron, 49 (6), 805–813.

Rutishauser, U., Ye, S., Koroma, M., Tudusciuc, O., Ross, I. B., Chung, J. M., & Mamelak, A. N. (2015). Representation of retrieval confidence by single neurons in the human medial temporal lobe. Nature neuroscience, 18 (7), 1041–1050.

Salz, D. M., Tiganj, Z., Khasnabish, S., Kohley, A., Sheehan, D., Howard, M. W., & Eichenbaum, H. (2016). Time cells in hippocampal area CA3. Journal of Neuroscience, 36, 7476–7484.

Schacter, D. L., Addis, D. R., & Buckner, R. L. (2007). Remembering the past to imagine the future: the prospective brain. Nature Reviews, Neuroscience, 8 (9), 657–661.

Schwartz, G., Howard, M. W., Jing, B., & Kahana, M. J. (2005). Shadows of the past: Temporal retrieval effects in recognition memory. Psychological Science, 16 (11), 898–904.

Sederberg, P. B., Howard, M. W., & Kahana, M. J. (2008). A context-based theory of recency and contiguity in free recall. Psychological Review, 115, 893–912.

Sederberg, P. B., Miller, J. F., Howard, M. W., & Kahana, M. J. (2010). The temporal contiguity effect predicts episodic memory performance. Memory & Cognition, 38, 689–699.

Singh, I., Oliva, A., & Howard, M. W. (2017). Visual memories are stored along a logarithmically-compressed representation of the past. Psychological Science.

Slotnick, S. D., & Dodson, C. S. (2005). Support for a continuous (single process) model of recognition memory and source memory. Memory & Cognition, 33, 151–170.

Squire, L. R., Wixted, J. T., & Clark, R. E. (2007). Recognition memory and the medial temporal lobe: a new perspective. Nature Reviews, Neuroscience, 8 (11), 872–83.

Staresina, B. P., Fell, J., Dunn, J. C., Axmacher, N., & Henson, R. N. (2013). Using state-trace analysis to dissociate the functions of the human hippocampus and perirhinal cortex in recognition memory. Proceedings of the National Academy of Sciences, 110 (8), 3119–24. doi:10.1073/pnas.1215710110

Teyler, T. J., & DiScenna, P. (1985). The role of hippocampus in memory: a hypothesis. Neuroscience and Biobehavioral Reviews, 9 (3), 377–89.

Teyler, T. J., & DiScenna, P. (1986). The hippocampal memory indexing theory. Behavioral Neuroscience, 100 (2), 147–54.

Tulving, E. (1972). Episodic and semantic memory. In E. Tulving & W. Donaldson (Eds.), Organization of memory. (p. 381–403). New York: Adademic Press.

Unsworth, N. (2008). Exploring the retrieval dynamics of delayed and final free recall: Further evidence for temporal-contextual search. Journal of Memory and Language, 59, 223–236.

Wixted, J. T. (2007). Dual-process theory and signal-detection theory of recognition memory. Psychological Review, 114 (1), 152–76.

Yaffe, R. B., Kerr, M. S. D., Damera, S., Sarma, S. V., Inati, S. K., & Zaghloul, K. A. (2014). Reinstatement of distributed cortical oscillations occurs with precise spatiotemporal dynamics during successful memory retrieval. Proceedings of the National Academy of Sciences, 111 (52), 18727–32. doi:10.1073/pnas.1417017112

Yonelinas, A. P. (2002). The nature of recollection and familiarity: A review of 30 years of research. Journal of Memory and Language, 46 (3), 441–517.

Yonelinas, A. P., Kroll, N. E., Quamme, J. R., Lazzara, M. M., Sauvé, M. J., Widaman, K. F., & Knight, R. T. (2002). Effects of extensive temporal lobe damage or mild hypoxia on recollection and familiarity. Nature Neuroscience, 5 (11), 1236–41.

